# The epithelial-specific ER stress sensor IRE1β enables host-microbiota crosstalk to affect colon goblet cell development

**DOI:** 10.1101/2021.07.28.453864

**Authors:** Michael J. Grey, Heidi De Luca, Doyle V. Ward, Irini A. M. Kreulen, Sage E. Foley, Jay R. Thiagarajah, Beth A. McCormick, Jerrold R. Turner, Wayne I. Lencer

## Abstract

Epithelial cells lining mucosal surfaces of the gastrointestinal and respiratory tracts uniquely express IRE1β (*Ern2*), a paralogue of the most evolutionarily conserved endoplasmic reticulum stress sensor IRE1α. How IRE1β functions at the host-environment interface and why a second IRE1 paralogue evolved remain incompletely understood. Using conventionally raised and germ-free *Ern2*^-/-^ mice, we found that IRE1β was required for microbiota-induced goblet cell maturation and mucus barrier assembly in the colon. This occurred only after colonization of the alimentary tract with normal gut microflora, which induced IRE1β expression. IRE1β acted by splicing *Xbp1* mRNA to expand ER function and prevent ER stress in goblet cells. Although IRE1α can also splice *Xbp1* mRNA, it did not act redundantly to IRE1β in this context. By regulating assembly of the colon mucus layer, IRE1β further shaped the composition of the gut microbiota. Mice lacking IRE1β had a dysbiotic microbial community that failed to induce goblet cell development when transferred into germ-free wild type mice. These results show that IRE1β evolved at mucosal surfaces to mediate crosstalk between gut microbes and the colonic epithelium required for normal homeostasis and host defense.

## INTRODUCTION

Epithelial cells lining mucosal surfaces of the gastrointestinal and respiratory tracts uniquely express two paralogues of the most evolutionarily conserved endoplasmic reticulum (ER) stress sensor IRE1. While all cell types express the essential stress sensor IRE1α, which mediates one arm of the unfolded protein response to ER stress, only mucosal epithelial cells at these sites express IRE1β (1). Why cells lining mucosal surfaces require a second IRE1 paralogue and how IRE1β contributes to the management of ER stress at the host-environment interface, however, remain incompletely understood.

Several lines of evidence point to a role for IRE1β in goblet cells, which are specialized exocrine cells that produce and secrete the mucin glycoproteins. In mice, expression of the *Ern2* gene, which codes for IRE1β, is highly enriched in goblet cells of the small intestine (2), colon (3), and respiratory tract (4). Deletion of *Ern2* reduces the number of MUC2^+^ goblet cells in the ileum (where MUC2 is the predominant mucin expressed in the intestine) (5) and induces the accumulation of misfolded MUC2 precursors (and ER stress) in secretory progenitor cells of the colon (6). These goblet cell phenotypes, however, are not present in mice with intestine specific deletion of IRE1α (encoded by *Ern1* gene), implicating specificity for IRE1β function (5, 6). Mucin glycoproteins produced by goblet cells assemble into the extracellular mucus layers lining mucosal surfaces. These mucus layers provide the initial interface guarding the adjacent epithelium from environmental factors, including the high density of microbes colonizing the gut (for recent review, see (7)). Notably, in the colon, the colonizing gut microbes contribute to the regulation of the mucus barrier (8, 9), and the composition of the mucus barrier shapes the colonizing microbial communities by providing a niche for growth and attachment (10). Although IRE1β contributes to goblet cell function, it is unknown how IRE1β contributes to the complex interface between microbes, the epithelium, and the mucus layers.

IRE1β, like IRE1α, is an ER transmembrane protein with a lumenal stress sensing domain and cytosolic kinase and endonuclease effector domains. The main signaling outputs for both proteins originate from the endonuclease domain, which enzymatically cleaves *Xbp1* mRNA in the cytosol to produce a spliced transcript coding for the transcription factor XBP1 (11, 12). The endonuclease domain can also degrade certain ER-targeted mRNAs to affect the cellular proteome through a process termed Regulated IRE1 Dependent Decay of mRNA (or RIDD) (13, 14). Both endonuclease activities may play a role in how IRE1β regulates mucin biosynthesis (6, 15). But, despite the high degree of sequence homology, IRE1β functions distinctly from IRE1α in that it has weaker endonuclease activity, responds only marginally to ER stress stimuli, and acts as a dominant-negative suppressor of stress-induced IRE1α signaling (16). These activities are partially explained by a non-conserved amino acid substitution in the kinase domain of IRE1β that impairs autophosphorylation, oligomerization, and thus stress-induced activation. We also note again the functions of IRE1β cannot simply be redundant to IRE1α, as the defects in goblet cell numbers and mucin biosynthesis found in *Ern2*^-/-^ mice are not found in mice with intestine-specific deletion of *Ern1* (5, 6), IRE1β but not IRE1α expression is enriched in goblet cells lining mucosal surfaces, and IRE1β expression is not found in other highly secretory cell types of other tissues that require IRE1α (e.g. pancreatic acinar and hematopoietic plasma cells).

Here, we used conventionally raised and germ free *Ern2*^-/-^ mice to elucidate the function of IRE1β in the intestinal mucosa. We found that IRE1β enables goblet cell maturation, mucin secretion, and mucus barrier assembly. Remarkably, expression of IRE1β and its functions in the colon were dictated by the gut microbiota, and IRE1β in turn, shaped the structure of the gut microbial communities. These results implicate IRE1β in mediating host-microbiota crosstalk that is critical for the development of the mucus barrier and host defense.

## RESULTS

### IRE1β enables goblet cell development in response to the gut microbiota

To determine how IRE1β contributes to mucosal homeostasis, we first compared morphology and genome-wide mRNA expression in colon crypt epithelial cells from wild type (WT) and *Ern2*^-/-^ mice. Under conventionally raised conditions (CONV), the colon epithelium of CONV-*Ern2*^-/-^ mice compared to CONV-WT littermate controls had significantly fewer goblet cells with smaller mucus vacuoles, as assessed by Alcian blue (AB) (Fig. 1A; Supporting Data (S.D.) 1A), periodic acid Schiff, and anti-MUC2 antibody staining as well as an expanded proliferative zone and elongated crypts (Supp. Fig. S1). Colon crypts from CONV-*Ern2*^-/-^ mice also had reduced mRNA expression for goblet cell signature genes (Fig. 1B; TableS1, S.D. 1B), including goblet cell transcription factors (*Atoh1*, *Spdef*), products (*Muc2*, *Retnlb*, *Itln1*), and genes generally related to biosynthetic secretory compartments that typify goblet cell function (Fig. 1B and 1C). Remarkably, and unlike in CONV-WT mice, treatment of CONV-*Ern2*^-/-^ mice with the γ-secretase inhibitor dibenzazepine (DBZ) to block Notch signaling and induce goblet cell differentiation (17) failed to increase the number of AB^+^ cells per crypt or upregulate goblet cell signature genes (Fig. 1D; Supp. Fig. S1; S.D. 1D). These data show that IRE1β is required for normal goblet cell development and implicate an IRE1β-dependent step downstream of niche factors that normally signal for differentiation and maturation of goblet cells in the colon.

**Figure 1.**
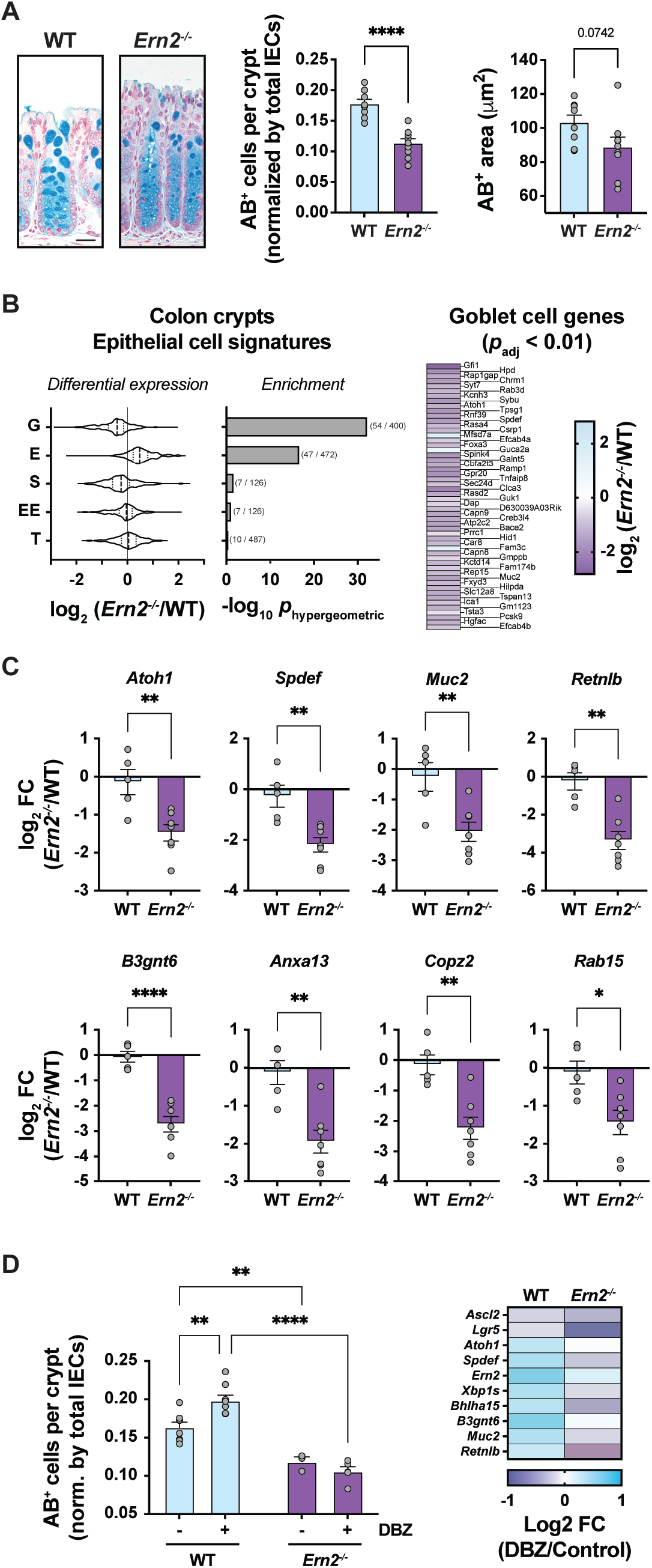
The colon epithelium of *Ern2*^-/-^ mice has fewer goblet cells under conventionally raised conditions. (A) Representative images of AB-stained sections of distal colon from CONV-WT and CONV-*Ern2*^-/-^ mice. Bar graphs show AB^+^ cells per crypt (normalized by the total number of crypt epithelial cells), and the AB-stained theca area of goblet cells measured in well-oriented crypts from the distal colon of CONV-WT and CONV-*Ern2*^-/-^ littermates. Symbols represent average values for individual mice (WT, n = 8; *Ern2*^-/-^, n = 9) from 2 independent cohorts. Bars represent mean ± SEM. Mean values were compared by unpaired t-test. The scale bar is equal to 50 μm and applies to both images. (B) Violin plot showing the distribution of relative mRNA expression of epithelial cell signature genes in colon crypts from CONV-*Ern2*^-/-^ mice compared to crypts from CONV-WT littermates (G = Goblet, E = Enterocyte, S = Stem, EE = Enteroendocrine, and T = Tuft). Heat map shows mean log2 fold change for differentially expressed goblet cell signature genes (n = 3 mice per group). (C) Bar graphs show relative mRNA expression of goblet cell signature genes in colon crypts measured by qPCR from an independent cohort of mice. Symbols represent individual mice and bars represent mean ± SEM; mean values were compared by unpaired t-test. (D) Bar graph shows the number of AB^+^ cells in upper half of well-oriented crypts (normalized by total number of crypt epithelial cells) for mice treated with or without the γ-secretase inhibitor DBZ. Symbols represent the average value an individual mouse and bars represent mean ± SEM. Mean values were compared by two-way ANOVA (n = 4-8 mice per group). Heat map shows differential expression (log2 [DBZ/Control]) of mRNA for goblet cell-related genes in colon epithelial cells following DBZ treatment as measured by qPCR.

Gut microbes are well known to affect goblet cell differentiation and the mucus layer that overlies the colonic epithelium. To investigate whether this depends on IRE1β, we rederived WT and *Ern2*^-/-^ mice under germ free (GF) conditions. As expected, we found that CONV-WT mice have significantly more goblet cells per crypt compared to germ free wild type (GF-WT) mice (Fig. 2A, light blue bars; S.D. 2A); and they had increased mRNA expression of goblet cell signature genes, including *Ern2* (Fig. 2B, 2C; TableS2 and S.D. 2B). Also as predicted, colonization of GF-WT mice with stool microbiota from CONV-WT donor mice restored the number of goblet cells per crypt (Fig. 2A) and restored the differential expression of goblet cell genes to levels normally found in CONV-WT mice (Fig. 2D; S.D. 2D, TableS3). All of this failed to occur, however, in *Ern2*^-/-^ mice. The presence of gut microbes in CONV-*Ern2*^-/-^ mice or following microbial colonization of GF-*Ern2*^-/-^ mice with microbes from CONV-WT donor mice had no detectable effect on the number of AB^+^ goblet cells per crypt (Fig. 2A, compare purple bars) or on goblet cell gene expression (Fig. 2B, 2C, and 2E; TableS4, TableS5, S.D. 2B, and S.D. 2E). We also found that GF-WT and GF-*Ern2*^-/-^ mice exhibited nearly identical goblet cell histology (Fig. 2A) and epithelial cell type specific gene expression patterns (Fig. 2F; Table S6, S.D. 2F). Thus, IRE1β is required for goblet cell development induced by the presence of gut microbes, implicating a relationship between IRE1β and the gut microbiota in shaping the mucosal environment of the mouse colon.

**Figure 2.**
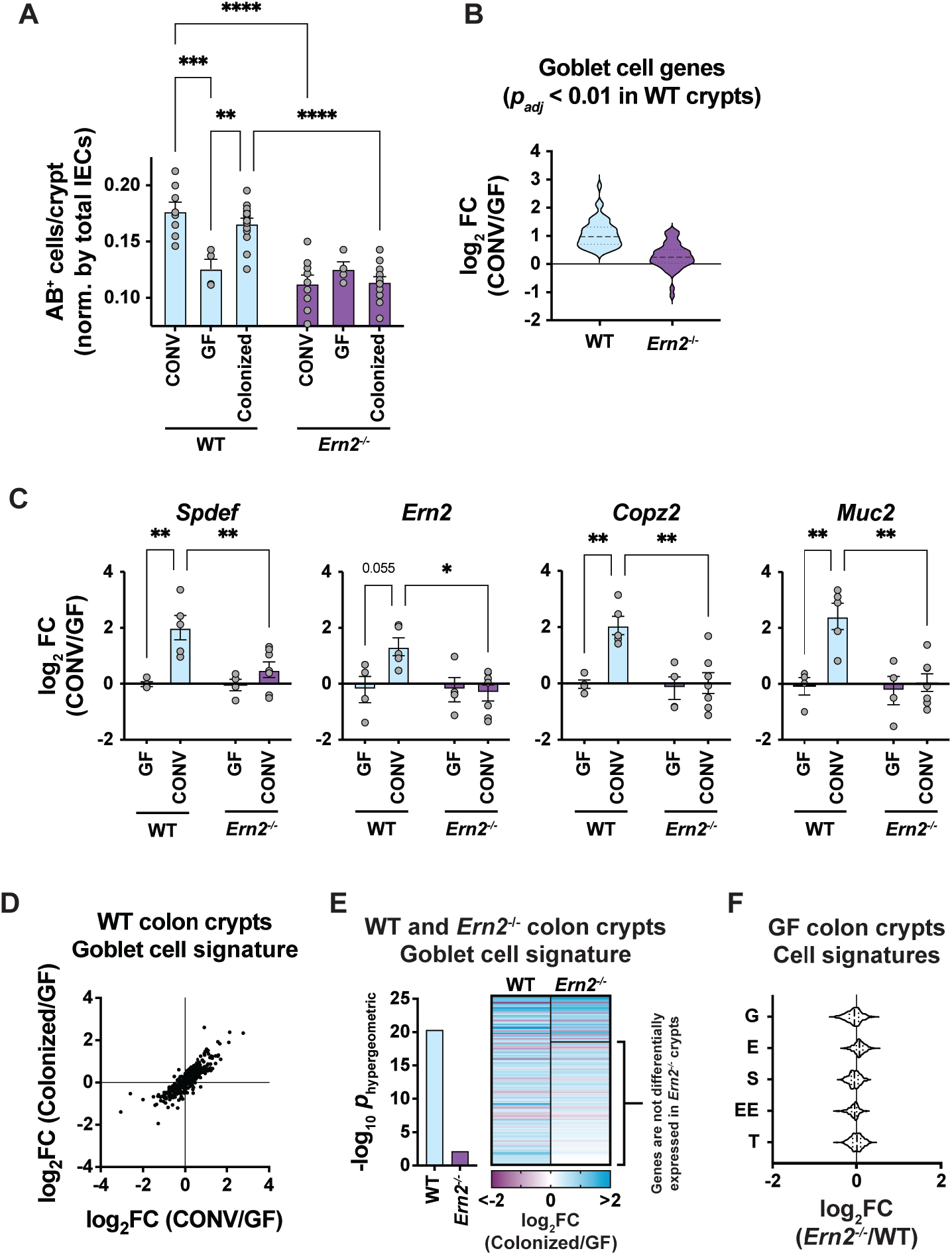
Gut microbes induce colon goblet cell development in an IRE1β-dependent manner. (A) Bar graph shows the number of AB^+^ cells in the upper half of well-oriented crypts in the distal colon of WT and *Ern2*^-/-^ under conventionally raised conditions (CONV), germ free conditions (GF), and GF followed by colonization with gut microbes from CONV-WT donor mice (COLONIZED). Symbols represent the average value for an individual animal and bars represent mean ± SEM. Mean values were compared by two-way ANOVA; mean values for GF-WT, CONV-*Ern2*^-/-^, GF-*Ern2*^-/-^, and COLONIZED-*Ern2*^-/-^ are not significantly different. Data for CONV mice is from Fig. 1A. (B) Violin plot showing relative mRNA expression for goblet cell signature genes that were upregulated in CONV-WT mice compared to GF-WT mice. Relative expression for those same genes is plotted for *Ern2*^-/-^ mice. (C) Bar graphs show relative mRNA expression (log_2_ [CONV/GF]) in colon crypts from WT and *Ern2*^-/-^ mice for select genes measured by qPCR. Symbols represent individual mice and bars represent mean ± SEM. Mean values were compared by two-way ANOVA. (D) Scatter plot compares differential expression of all goblet cell signature genes for CONV-WT versus GF-WT compared to COLONIZED-WT versus GF-WT. (E) Bar graph shows enrichment of differentially expressed genes from the goblet cell signature for COLONIZED-WT and COLONIZED-*Ern2*^-/-^ mice compared to GF controls. The heat map shows the relative expression of the differentially expressed genes. A subset of the genes that are differentially expressed in WT mice are not differentially expressed (*p_adj_* > 0.01) in *Ern2*^-/-^ mice as indicated. (F) Violin plot showing the distribution of relative mRNA expression of epithelial cell signature genes from colon crypts of GF-*Ern2*^-/-^ mice compared to GF-WT mice (G = Goblet, E = Enterocyte, S = Stem, EE = Enteroendocrine, and T = Tuft).

### IRE1β-mediated Xbp1 splicing and XBP1 are required to expand ER function and mediate goblet cell maturation

Like IRE1α, IRE1β could affect epithelial gene expression by splicing *Xbp1* mRNA to produce the transcription factor XBP1 and/or by selectively degrading mRNA for other gene products via RIDD. To discern between these two mechanisms, we analyzed gene expression profiles for colon crypt epithelial cells from WT and *Ern2*^-/-^ mice. Colon crypts from CONV-WT and COLONIZED-WT mice had significant enrichment of upregulated genes associated with the transcription factor XBP1 compared to GF-WT mice (CONV vs GF, gProfiler TF:M01770_1, −log_10_ *p_adj_* = 15.48; COLONIZED vs GF, −log_10_ *p_adj_* = 15.12). This microbial-induced enrichment of XBP1-associated genes, however, was not apparent in CONV-*Ern2*^-/-^ or COLONIZED-*Ern2*^-/-^ mice. Of the 132 goblet cell signature genes differentially expressed in COLONIZED-WT mice, 96 of them were not differentially expressed in COLONIZED-*Ern2*^-/-^ mice compared to GF-*Ern2*^-/-^ mice (Fig. 2E; S.D. 2E). These genes were significantly associated with XBP1 (-log_10_ *p_adj_* = 9.44) and, as expected for IRE1-XBP1 signaling (18), ER function, protein processing in the ER, and ER-to-Golgi transport (Table 1; S.D. 2E). These results implicate XBP1 as a downstream effector of IRE1β in mediating microbial-induced gene expression and goblet cell maturation.

**Table 1.** Functional analysis of goblet cell genes that are differentially expressed in WT COLONIZED crypts but not Ern2^-/-^ COLONIZED crypts.

| **GO:BP** | **-log<sub>10</sub>(*p*<sub>adj</sub>)** |
| --- | --- |
| Response to ER stress | 4.71 |
| Protein Transport | 4.71 |
| ER to Golgi vesicle-mediated transport | 4.18 |
| Protein localization | 3.43 |
| Protein N-linked glycosylation | 3.20 |
| **GO:CC** |  |
| Endoplasmic reticulum | 13.73 |
| Golgi | 7.05 |
| Oligosaccharyltransferase complex | 6.34 |
| COPII-coated ER to Golgi transport | 5.87 |
| **KEGG** |  |
| Protein processing in the ER | 16.34 |
| **REAC** |  |
| N-linked glycosylation | 4.20 |
| ER to Golgi anterograde transport | 1.40 |
| **TF** |  |
| XBP1 | 9.44 |

To test if XBP1 affects goblet cell maturation, we analyzed the colonic epithelium of conventionally-raised mice with intestine-specific deletion of *Xbp1* (*Xbp1*^fl/fl^;*Vil-Cre*^+^). We found that CONV-*Xbp1*^fl/fl^;*Vil-Cre*^+^ mice closely phenocopied the CONV-*Ern2*^-/-^ mice. They had fewer AB^+^ goblet cells per crypt (Fig. 3A; S.D. 3A) and reduced expression of goblet cell signature genes compared to *Xbp1*^fl/fl^;*Vil-Cre*^-^ littermate controls (Fig. 3B; Supp. Fig. S3A). Thus, XBP1, like IRE1β, is required for goblet cell development in the colon. The result is consistent with earlier studies in mice that showed deletion of *Xbp1* impaired differentiation of secretory lineages along the gastrointestinal tract, including goblet cells in the small intestine (19, 20); and it further implicated XBP1 as the downstream effector of IRE1β.

**Figure 3.**
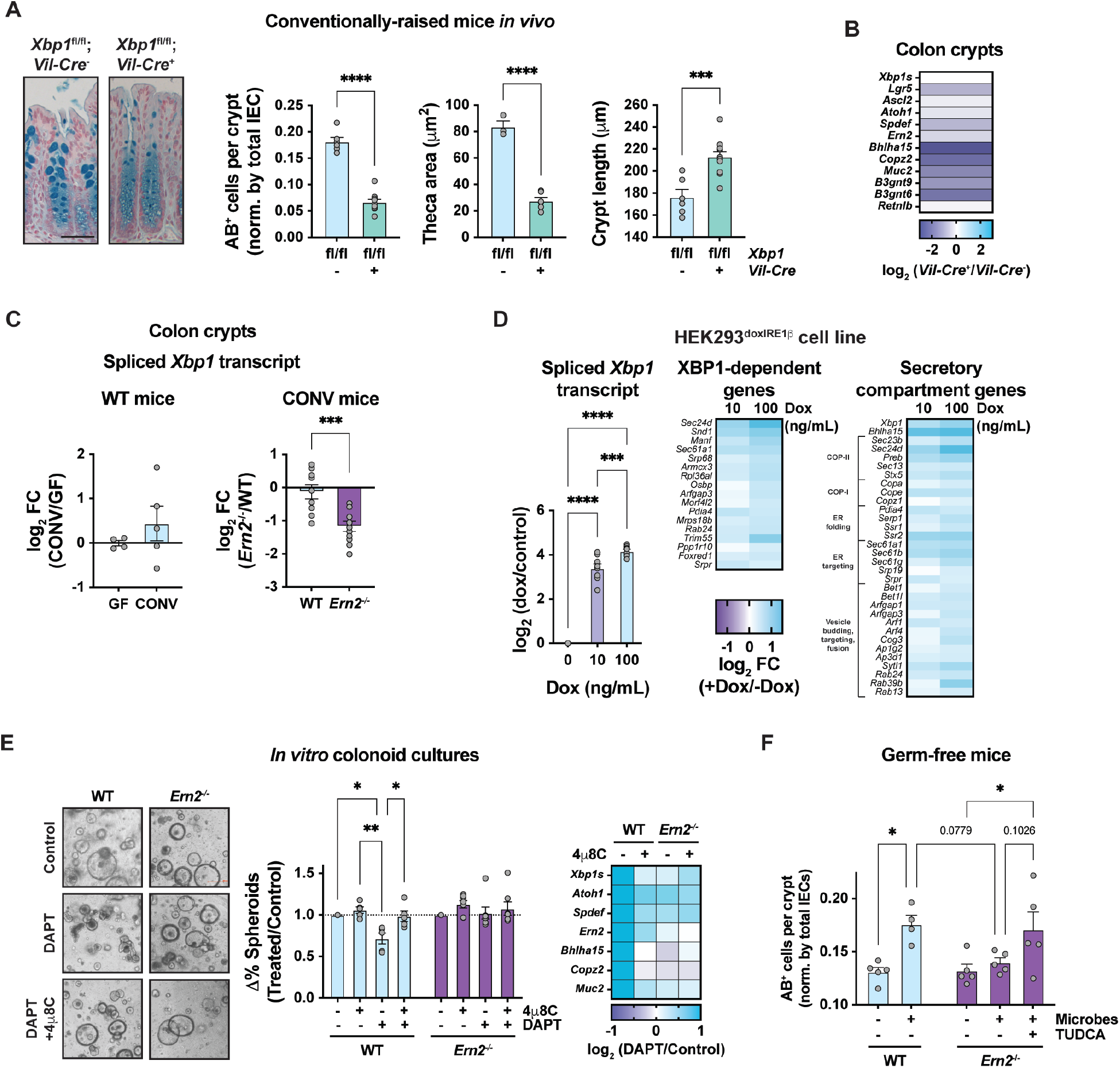
IRE1β-mediated *Xbp1* splicing and XBP1 are required to expand ER function and mediate goblet cell maturation. (A) Representative images of AB-stained sections of distal colon from *Xbp1*^fl/fl^;*Vil-Cre*^-^ and *Xbp1*^fl/fl^;*Vil-Cre*^+^ littermates. Bar graphs show the number of AB^+^ cells, the AB-stained goblet cell theca area, and crypt lengths measured in well-oriented crypts. Symbols represent individual mice from 2 independent cohorts (*Xbp1*^fl/fl^;*Vil-Cre*^-^, n = 6; *Xbp1*^fl/fl^;*Vil-Cre*^+^, n = 12) and bars represent mean ± SEM. Mean values were compared by unpaired t-test. The scale bar is 50 μm and applies to both images. (B) Heat map shows relative mRNA expression (log_2_ [*Vil-Cre*^+^/*Vil-Cre*^-^]) of goblet cell marker genes in colon crypt epithelial cells as measured by qPCR. (C) Bar graph shows relative expression of spliced *Xbp1* transcript in colon crypts from (left panel) GF-WT (n = 4) and CONV-WT mice (n = 6) and (right panel) CONV-WT (n = 9) and CONV-*Ern2*^-/-^ (n = 11) mice. Symbols represent individual mice and bars represent mean ± SEM. Mean values were compared by unpaired t-test. (D) Bar graph shows the fold change in relative expression of spliced *Xbp1* transcript in HEK293^doxIRE1β^ cells (16) treated with doxycycline at indicated concentration to induce IRE1β transgene expression. Symbols represent independent experiments and bars represent mean ± SEM. Mean values compared by one-way ANOVA (n = 9). Heat maps show mean values of log_2_ fold change in expression (relative to uninduced control cells) of genes associated with XBP1 and genes associated with secretory compartment function (n = 3 per group). Data in left panel are reproduced from Grey et al. (16). (E) Representative images of mouse colonoids treated with the γ-secretase inhibitor DAPT in the presence or absence of the IRE1 inhibitor 4μ8C. Differentiation status was assayed by scoring spheroid versus non-spheroid morphology. Bar graph shows the change in the percentage of colonoids with spheroid morphology relative to untreated controls for a given colonoid line within an experiment. Symbols represent independent experiments with at least 2 independent colonoid lines (WT, n = 5; *Ern2*^-/-^, n = 7). Mean values within genotype were compared by two-way ANOVA. Heat map shows relative mRNA expression (log_2_ [DAPT/Control]) of goblet cell marker genes measured by qPCR from a single experiment. (F) Bar graph shows the number of AB^+^ cells in upper half of well-defined crypts (normalized by total IECs) following colonization of GF-*Ern2*^-/-^ mice with a microbiota from CONV-WT donor mice in the presence or absence of TUDCA. Symbols represent the average value for an individual mouse and bars represent mean ± SEM (WT: GF/colonized, n = 5/4; *Ern2*^-/-^: GF/colonized/colonized+TUDCA, n = 5/5/5). Mean values were compared by one-way ANOVA.

This interpretation is supported by direct measurement of spliced *Xbp1* mRNA, which is needed for XBP1 translation (11, 12). Spliced *Xbp1* mRNA was induced by gut microbes in CONV-WT mice compared to GF-WT mice (Fig. 3C, left panel). However, colon crypt epithelial cells from CONV-*Ern2*^-/-^ mice had significantly less spliced *Xbp1* mRNA compared to CONV-WT mice (Fig 3C, right panel). Furthermore, *in vitro* studies showed that expression of IRE1β in HEK293 cells resulted in *Xbp1* mRNA splicing (Fig. 3D, left panel; Supp. Fig. S3B, and (16)), together with upregulation of XBP1-dependent genes and genes associated with ER function, protein folding, and ER-to-Golgi vesicle trafficking (Fig. 3D, middle and right panels; S.D. 3D, TableS8, TableS9). The results of these studies are consistent with IRE1β-mediated *Xbp1* splicing as mechanism of action for goblet cell maturation. To test if IRE1β enzymatic activity (and indirectly *Xbp1* splicing) is needed for goblet cell differentiation, we used an *in vitro* colonoid differentiation assay. Treatment of WT colonoid cultures with the γ-secretase inhibitor DAPT to induce goblet cell differentiation decreased the percentage of colonoids with a spheroid morphology (Fig. 3E, left and middle panels; S.D. 3E) and as expected, increased expression of goblet cell marker genes (Fig. 3E right panel, first column). *Ern2*^-/-^ colonoids, however, did not respond to DAPT (Fig. 3E middle panel and right panel, third column). Thus, as was the case *in vivo* with DBZ (Fig. 1D), IRE1α was not sufficient to mediate goblet cell differentiation in colonoid culture. Furthermore, the endonuclease activity of IRE1β was required, as treatment of WT colonoids with the IRE1 endonuclease inhibitor 4μ8C (21) blocked DAPT-induced goblet cell differentiation (Fig. 3E middle panel fourth column, and right panel second column). Based on these studies, we conclude that goblet cell development in the mouse colon depends upon expression of IRE1β and its function in enzymatically splicing *Xbp1* mRNA. Both are induced by gut microbes, and both are specific for IRE1β as IRE1α is unable to fulfill this function.

As XBP1 normally expands ER function to enable proteostasis, depletion of XBP1 in *Ern2*^-/-^ mice may result in ER stress following colonization. Accordingly, we found that crypts from COLONIZED-*Ern2*^-/-^ mice had significant enrichment of genes associated with the response to ER stress (GO:0034976, −log_10_ *p_adj_* = 5.15), with *Derl3*, *Chac1*, and *Trib3* being the most upregulated genes in crypts of COLONIZED-*Ern2*^-/-^ mice compared to COLONIZED-WT mice (TableS7, S.D. 3F). COLONIZED-*Ern2*^-/-^ mice also had decreased expression of genes involved in translation (GO:0006412, −log_10_ *p_adj_* = 18.23) and ribosome function (GO:0003735, −log_10_ *p_adj_* = 34.79; GO:0042255, −log_10_ *p_adj_* = 7.86; KEGG:03010, −log_10_ *p_adj_* = 34.22), suggesting that *Ern2*^-/-^ crypt epithelial cells have abnormally slowed protein synthesis in response to ER stress following colonization (S.D. 3F). Treatment of COLONIZED-*Ern2*^-/-^ mice with the chemical chaperone tauroursodeoxycholic acid (TUDCA) (22, 23) to relieve ER stress enabled the induction of AB^+^ goblet cells to a similar extent as COLONIZED-WT mice (Fig. 3F). From these results, we conclude that IRE1β function is required for the maturation of goblet cells in response to gut microbes by expanding ER and secretory compartment function to prevent the accumulation of ER stress.

### Ern2^-/-^ mice have impaired mucus assembly and earlier onset of infectious colitis

A primary function of goblet cells is to secrete mucin glycoproteins that assemble into mucus layers protecting the epithelium. In the distal colon, this includes a dense inner mucus layer that is impenetrable to microbes and separates them from the epithelial cell surface (24). A second loosely attached outer mucus layer also forms in the colon, but this layer is colonized by microbes. As such, to determine if IRE1β and XBP1 affect mucus assembly, we analyzed the mucus layers in Carnoy’s-fixed section of CONV-WT, CONV-*Ern2*^-/-^, and CONV-*Xbp1*^fl/fl^;*Vil-Cre*^+^ mice. Consistent with the defect in goblet cell development, the distal colon of CONV-*Ern2*^-/-^ mice had impaired mucus assembly with a significantly thinner inner mucus layer compared to CONV-WT mice (Fig. 4A; S.D. 4A-B). The same perturbations were found in mice with intestine-specific deletion of *Xbp1* (Fig. 4B). In some instances, the inner mucus layer was absent, and gut microbes were abnormally located immediately adjacent to the surface epithelial cells (Fig. 4C, EUB338 in situ hybridization probe shown in red). As in earlier studies (25), we also found that formation of the inner mucus layer was microbe dependent (Fig. 4D), again linking IRE1β with goblet cell development induced by microbial colonization.

**Figure 4.**
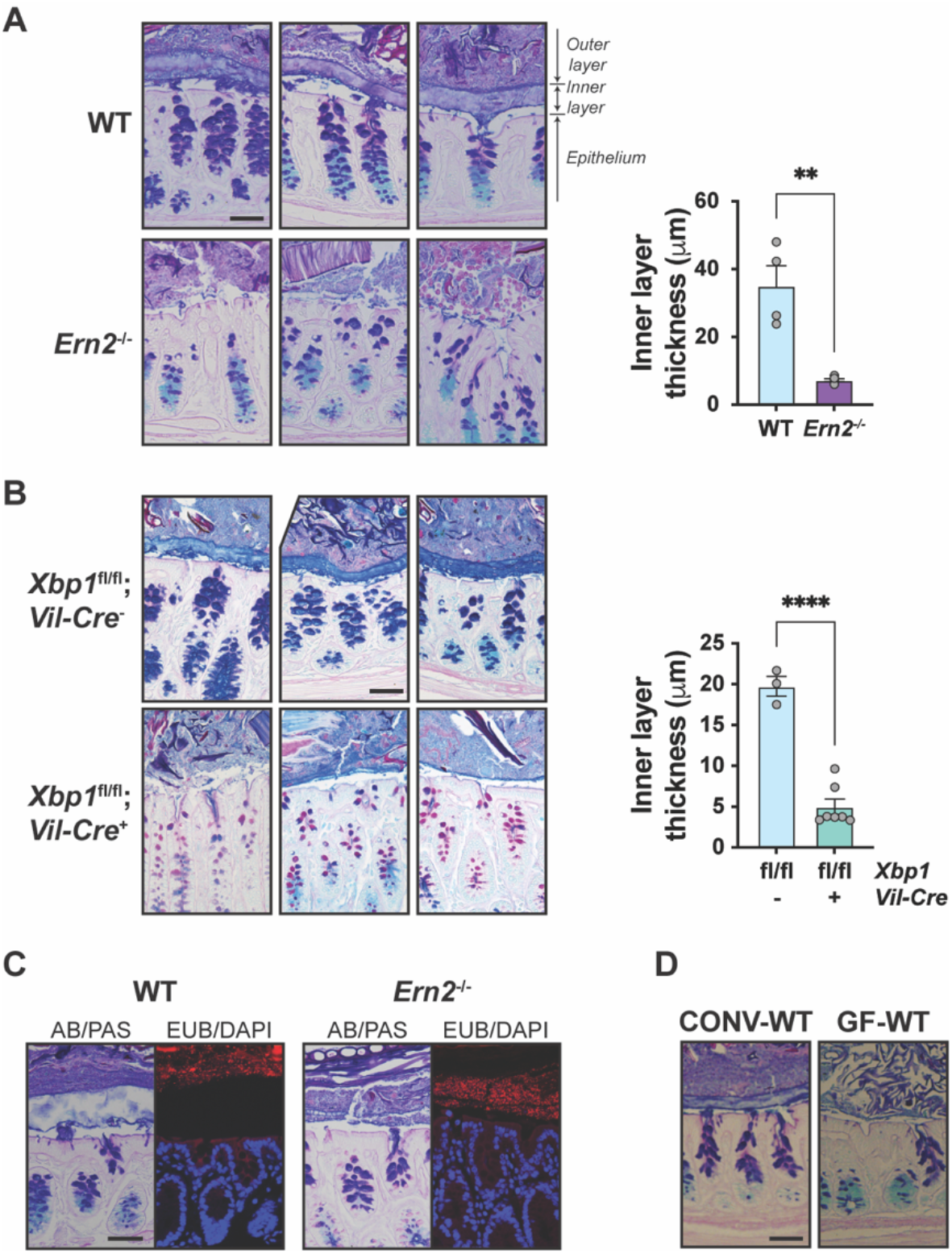
Impaired assembly of the colon mucus layer in *Ern2*^-/-^ mice. Representative images of the colon mucus layer assessed by AB/PAS staining of Carnoy’s-fixed tissue from (A) CONV-WT and CONV-*Ern2*^-/-^ mice and (B) *Xbp1*^fl/fl^;*Vil-Cre*^-^ and *Xbp1*^fl/fl^;*Vil-Cre*^+^ mice. Bar graphs show the thickness of the inner mucus layer from AB/PAS-stained sections. Symbols represent the average value of the distribution of measures around at least 2 full cross sections for individual mice (WT, n = 4; *Ern2*^-/-^, n = 5; *Xbp1*^fl/fl^;*Vil-Cre*^-^, n = 3; *Xbp1*^fl/fl^;*Vil-Cre*^+^, n = 7) and bars represent mean ± SEM. Mean values were compared by unpaired t-test. (C) Representative images of Carnoy’s-fixed colon tissue from CONV-WT and CONV-*Ern2*^-/-^ mice stained with AB/PAS to detect the colon mucus layer or stained with the 16S rRNA in situ hybridization probe EUB338 (red) to detect lumenal bacteria. Nuclei were stained with DAPI (blue). (D) Representative images of Carnoy’s fixed tissue from CONV-WT and GF-WT mice stained with AB/PAS to detect the colon mucus layer. The scale bars in (A) - (D) are equal to 50 μm and apply to all images shown.

We took a second approach to test functionally for IRE1β-dependent assembly of an intact mucus layer using the mouse pathogen *Citrobacter rodentium* (26, 27). Here, we found that conventionally raised *Ern2*^-/-^ mice had an earlier onset of *C. rodentium* infection as assessed by pathogen burden (CFUs cultured from stool) and day of onset (Fig. 5A; S.D. 5A). At 8 days post-infection, WT and *Ern2*^-/-^ mice both had significantly elevated levels of *C. rodentium* in stool compared to uninfected controls, but *Ern2*^-/-^ mice had on average ∼10-fold higher CFUs per gram of stool (Fig. 5B, left panel). This was associated with more adherent *C. rodentium* cultured from colon tissue (Fig. 5B, right panel) and a greater extent of histologic damage in *Ern2*^-/-^ mice compared to WT cagemate controls (Fig. 5C-E; S.D. 5B-D). Epithelial features of goblet cell depletion, mucosal hyperplasia, crypt cell apoptosis, and erosion were more pronounced in *Ern2*^-/-^ mice (Fig. 5D). After peak levels of infection were attained, WT and *Ern2*^-/-^ mice had similar degrees of pathogen burden and histologic damage (e.g. 13 days post-infection, Fig. 5F,G). But, despite the earlier onset of disease, *Ern2*^-/-^ mice were more efficient at clearing the infection (Fig. 5F); and they recovered more quickly as assessed by tissue histology (Fig. 5G; S.D. 5G) – consistent with previous reports that mucus delays *C. rodentium* clearance and resolution (28).

**Figure 5.**
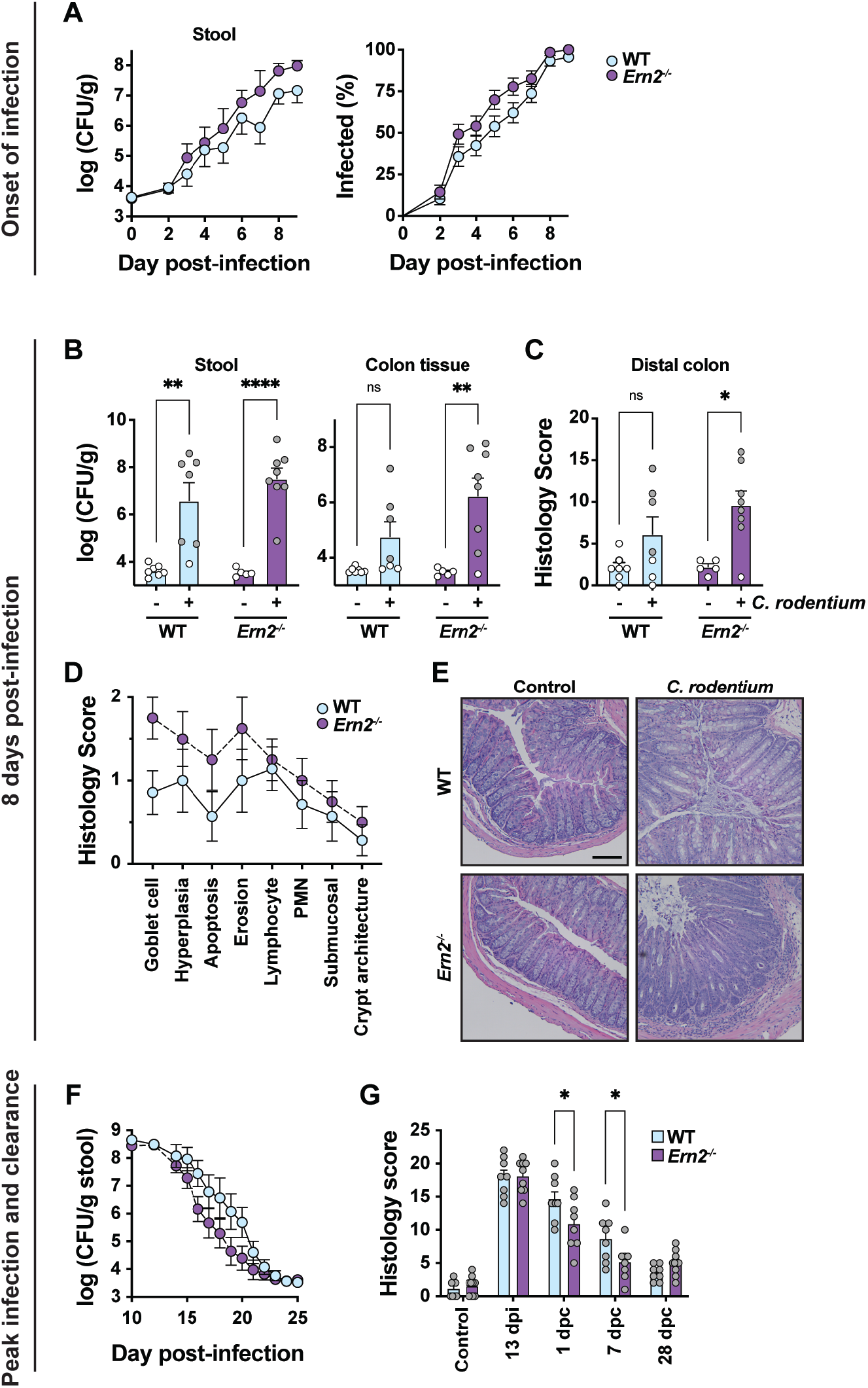
*Ern2*^-/-^ mice have earlier onset of *C. rodentium* infection. (A) Time courses represent the onset of infection monitored (left panel) by enumerating *C. rodentium* colony forming units (CFUs) in stool of infected mice and (right panel) by comparing the percentage of mice that tested positive over the first 9 days of infection. The time courses represent the average measured in 47 WT and 38 *Ern2*^-/-^ mice from 5 independent experiments (the number of mice in each time point varies). For the CFU time course, the value at time zero is equal to the minimum CFUs that can be detected in the assay. (B) Bar graphs show *C. rodentium* CFUs cultured from (left panel) stool and (right panel) colon tissue from control and infected mice at 8 days post-infection (WT, n = 7/7 control/infected; *Ern2*^-/-^, n = 5/8 control/infected). Symbols represent individual mice and bars represent mean ± SEM. Mice that did not have any CFUs detected are shown in white and are assigned the minimum CFUs that can be detected in the assay for a given stool/tissue sample. Mean values were compared by two-way ANOVA. (C) Bar graph shows histology scores reflective of epithelial damage and inflammation in distal colon of control and infected mice at 8 days post-infection. Symbols represent individual mice and those in white correspond to mice that did not have *C. rodentium* CFUs detected in colon tissue from (B). (D) Plot shows the average value for each histologic parameter, including goblet cell depletion, mucosal hyperplasia, crypt apoptosis, epithelial erosion, lympocytic infiltrate, neutrophilic infiltrate, submucosal inflammation and edema, and crypt architectural distortion, for infected mice scored in (C). (E) Representative images of H&E-stained section of distal colon from control and infected mice at 8 days post-infection. The scale bar is equal to 100 μm and applies to all images. (F) Time course represents clearance of infection monitored by enumerating *C. rodentium* CFUs in stool of infected mice. Symbols represent mean ± SEM (WT, n = 14; *Ern2*^-/-^ n = 13). (G) Bar graphs show histology scores evaluated in H&E-stained sections of distal colon from uninfected (Control) mice and infected mice at 13 days post-infection, and 1-, 7-, and 28-days post-clearance (*C. rodentium* CFUs no longer detected in stool).

### IRE1β mediates a Microbiota—Epithelial—Mucus feedback loop that maintains mucosal homeostasis

The gut microbiota and the colonic mucus layer are linked via a feedback loop, where specific components of the gut microbiota are associated with development of the colon mucus layer (9, 25), and the mucus layer in turn selects for different bacterial species (10). As such, we hypothesized that the microbiota colonizing WT and *Ern2*^-/-^ mice might be different and may contribute to the *Ern2*^-/-^ phenotype. To test this, we first asked if the microbiota from *Ern2*^-/-^ mice could phenocopy the microbiota from WT mice by inducing goblet cell development when transferred into recipient GF-WT mice (as in Fig. 2A). We also tested this using CONV-WT recipient mice pretreated with antibiotics to deplete the gut microbiota. In both models, we found that colonization with stool microbiota from CONV-WT donor mice fully restored goblet cell numbers as expected (Fig. 6A and 6B; S.D. 6A, S.D. 6B). In contrast, the stool microbiota from CONV-*Ern2*^-/-^ donor mice failed to rescue the normal goblet cell phenotype in either recipient model (Fig. 6A and 6B). To document how the loss of IRE1β might have altered the gut microbiota in *Ern2*^-/-^ mice, we used metagenomic sequencing to compare the composition of stool microbes obtained from co-housed CONV-WT and CONV-*Ern2*^-/-^ mice. The analysis showed that even after co-housing, the microbiota from *Ern2*^-/-^ mice was less diverse than that of WT mice as indicated by the significant reduction in Shannon (taxa abundance) and Faith (phylogenetic diversity) measures of alpha diversity (Fig. 6C). We also found that the taxa of the two microbial communities diverged significantly as measured by beta diversity using weighted UniFrac (*p* = 0.012), Bray-Curtis dissimilarity (*p* = 0.005), or Jaccard distance (*p* = 0.007). We note in particular that the stools of *Ern2*^-/-^ mice showed a significant reduction in relative abundance of taxa belonging to the *Firmicutes* phylum overall, as well as species known to promote mucus barrier function (e.g. *Lactobacillus reuteri* (29), Fig. 6D; S.D. 6D). Thus, loss of IRE1β function results in a dysbiotic gut microbiota that is unable to promote goblet cell development when transferred into GF-WT mice.

**Figure 6.**
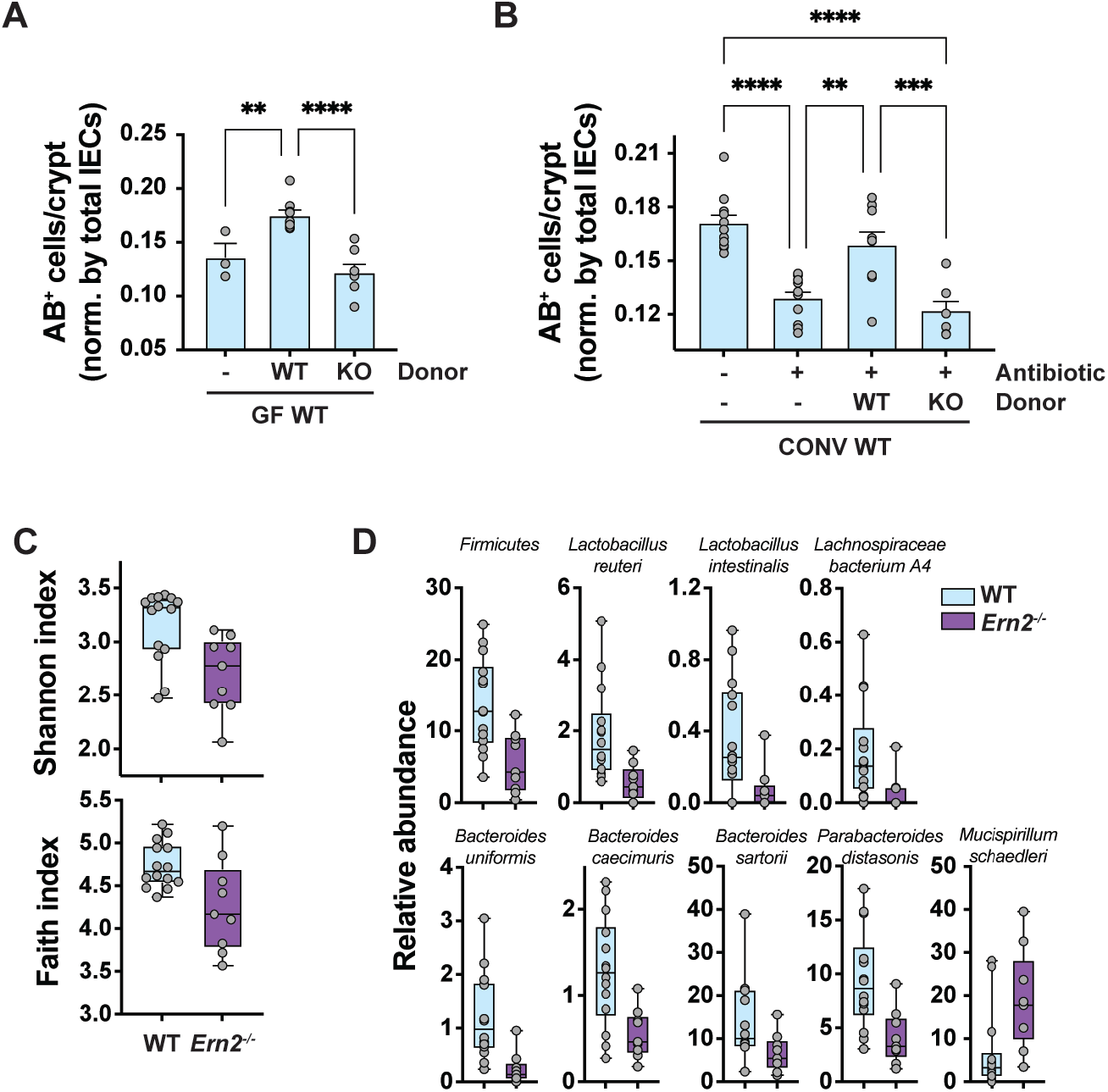
Epithelial goblet cell defect transfers with the *Ern2*^-/-^ microbiota. Bar graphs show number of AB^+^ cells per crypt in the upper half of well-oriented crypts in the distal colon of (A) GF-WT mice (n = 3) and GF-WT mice colonized with microbiota from WT (n = 9) or *Ern2*^-/-^ (n = 8) donor mice and (B) CONV-WT mice (n = 11), antibiotic-treated CONV-WT mice (n = 11), and antibiotic-treated CONV-WT mice that were recolonized by cohousing with WT (n = 9) or *Ern2*^-/-^ (n = 7) donor mice. Symbols represent the average value for an individual mouse and bars represent mean ± SEM. Mean values were compared by one-way ANOVA. (C) Box plots show alpha diversity indices for microbiota from WT and *Ern2*^-/-^ mice. Symbols represent values for an individual mouse and error bars represent min and max. (D) Box plots show relative abundance for indicated taxa that are significantly different between microbiota from CONV-WT and CONV-*Ern2*^-/-^ mice. Symbols represent relative abundance for an individual mouse. False discovery rate q values were calculated with MaAsLin2 (WT, n = 14; *Ern2*^-/-^, n = 9).

As a next step to understand what components of the gut microbiota modulate IRE1β expression and goblet cell development, we tested the short-chain fatty acid butyrate. *In vitro* treatment of polarized T84 cell monolayers with butyrate, but not acetate, increased *Ern2* mRNA expression as well as spliced *Xbp1* mRNA (Fig. 7A). The response to butyrate was replicated in LS174T cells (30), which also showed butyrate induced *Muc2* mRNA expression that typifies goblet cell differentiation (Fig. 7B). When tested *in vivo*, GF-WT mice given drinking water with butyrate had an increase in the number of colonic AB^+^ goblet cells to the level induced by colonization with a microbiota from CONV-WT donor mice. In contrast, there was no effect on goblet cell numbers when GF-*Ern2*^-/-^ mice were given butyrate in their drinking water (Fig. 7C). Thus, microbially-produced butyrate may link certain gut microbiota with *Ern2* expression and goblet cell development in the colon epithelium. Consistent with this interpretation, the metagenome of CONV-*Ern2*^-/-^ mice had reduced representation of butyrate kinase and phosphate butyryltransferase genes (Fig. 7D) – enzymes that catalyze final steps in butyrate production. We also found that stool from CONV-*Ern2*^-/-^ mice had a lower mole fraction of butyrate (with slight increases in acetate and propionate) in comparison to stool from CONV-WT mice (Fig. 7E). Thus, we conclude that butyrate is at least one component of the gut microbiota that regulates *Ern2* expression and goblet cell development in the colonic epithelium.

**Figure 7.**
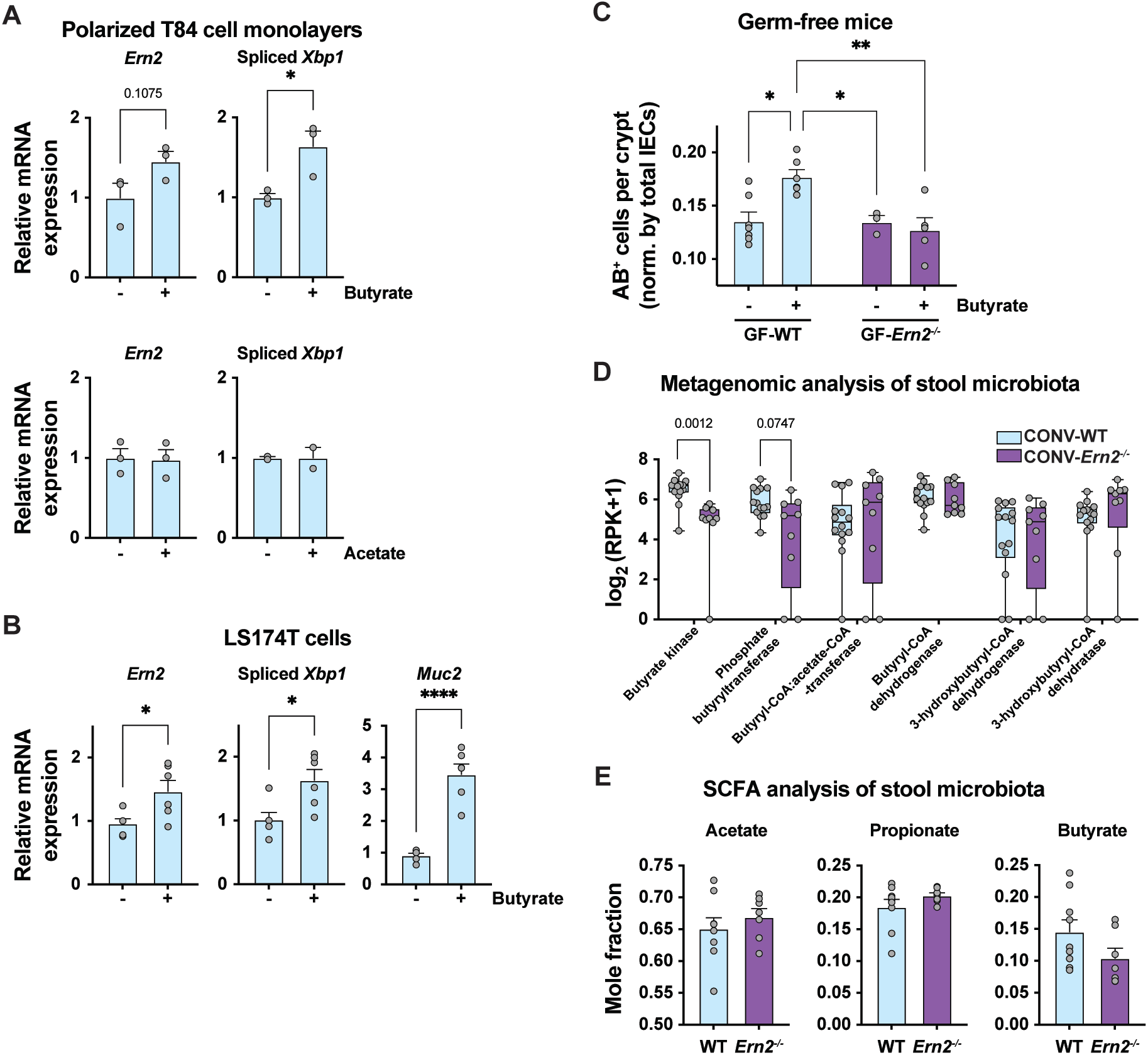
The short chain fatty acid (SCFA) butyrate links gut microbes with *Ern2* expression. Bar graphs show relative expression of *Ern2* and spliced *Xbp1* mRNA measured by qPCR in (A) polarized T84 cell monolayers (*n* = 3) and (B) LS174T cells (*n* = 6) treated with butyrate or acetate as indicated. Symbols represent independent experiments and bars represent mean ± SEM; mean values were compared by t-test (unpaired, two-tailed). (C) Bar graph shows the number of AB^+^ cells per crypt in the distal colon of GF-WT and GF-*Ern2*^-/-^ mice that received regular drinking water or drinking water supplemented with sodium butyrate. Symbols represent the average value for an individual mouse and bars represent mean ± SEM; mean values were compared by two-way ANOVA. (D) Box plots show whole genome sequence reads for enzymes in butyrate metabolism measured in stool from WT (*n* = 14) and *Ern2*^-/-^ (*n* = 9) mice. Symbols represent an individual mouse and bars represent the range. Mean values were compared using multiple t-tests. (E) Bar graphs show mole fraction of SCFAs measured in stool from WT (*n* = 9) and *Ern2*^-/-^ (*n* = 7) mice. Values are shown as the mole fraction of each relative to the total SCFAs measured in a given sample. Symbols represent an individual mouse and bars represent mean ± SEM.

## DISCUSSION

Our results define an essential role for the epithelial-specific ER stress sensor IRE1β in the crosstalk between gut microbiota and the colonic epithelium that drives goblet cell differentiation and development of an effective mucus barrier. *Ern2* gene transcription and IRE1β expression are indispensable to this process, and both are induced by gut microbial colonization. In turn the mucus barrier produced following IRE1β expression selects for a colonizing microbial community that maintains *Ern2*/IRE1β expression, and thus goblet cell differentiation and mucus barrier assembly. The ubiquitously expressed IRE1α paralogue does not function redundantly to IRE1β in this context. In mice lacking *Ern2* but expressing *Ern1*, goblet cells fail to differentiate in response to microbes, and the defective mucus barrier produced selects for a dysbiotic microbial community that cannot induce *Ern2* gene expression in WT animals. Thus, unlike IRE1α, the IRE1β paralogue enables a crosstalk between host and microbe that is essential for development and maintenance of mucosal tissues.

IRE1β appears to act by splicing *Xbp1* to affect goblet cell maturation and the mucus barrier. This is in line with a broader view of XBP1 function in development of multiple secretory lineages along the GI tract (5, 19, 20), including, as shown here, goblet cells populating the colon epithelium. Ribiero and coworkers also proposed that IRE1β acts via XBP1 in mediating allergen-induced development of mucus producing cells in the airway epithelium (15). We cannot, however, rule out contribution of IRE1β endonuclease activity via RIDD. Kohno and coworkers reported that IRE1β RIDD activity degrades *Muc2* mRNA as a mechanism to post-transcriptionally regulate MUC2 protein expression (and thus ER stress) (6). It is noteworthy, however, that we do not see accumulation of *Muc2* mRNA in colon crypt epithelial cells at steady state in CONV-*Ern2*^-/-^ mice or following colonization of GF-*Ern2*^-/-^ mice – as one might expect in colon tissues lacking IRE1β-mediated RIDD activity. Rather, crypts from *Ern2*^-/-^ mice tended to have decreased levels of *Muc2* mRNA compared to their WT counterparts. But we did not assay *Muc2* mRNA stability in our studies, and the decrease in *Muc2* mRNA levels we observe in crypts from *Ern2*^-/-^ mice likely reflects lower goblet cell numbers and their failure to fully differentiate. IRE1β function via RIDD remains a plausible and additional mechanism of action.

Different hypotheses have been proposed to explain why epithelial cells at mucosal surfaces uniquely express the two IRE1 paralogues. As we have noted before, the two IRE1 paralogues are found only in vertebrates where the *Ern2* gene likely arose after a whole genome duplication event that typifies vertebrate genomes (31). In one mechanism to retain the duplicated gene, a relaxed selective pressure on *Ern2* may have allowed for the accumulation of sequence variations that enabled the selection of new functions – a process called neofunctionalization. The *Ern2* genes in mammals have acquired the most pronounced sequence variation compared to their *Ern1* counterparts (31). This, we suggest, is a strong hint towards explaining the unique tissue distribution of *Ern2* gene expression. A defining feature of epithelial barriers in mammals, compared to lower vertebrates where *Ern1* and *Ern2* genes are more similar, and compared to invertebrates that have only a single IRE1 gene, is the evolution of a mucus-based rather than a chitin-based system of host-environment barrier immunity (31, 32). And this, as we show here, requires *Ern2*/IRE1β. Though other hypotheses have been proposed to explain why epithelial cells at mucosal surfaces may have evolved to express IRE1β (1, 5, 16, 33, 34), we propose the evolutionary shift to a mucus based system of mucosal immunity in mammals underlies its evolution, and at least one of its major functions.

Furthermore, although both IRE1α and IRE1β paralogues signal via their endonuclease activity in similar ways (*Xbp1* mRNA splicing and RIDD), we know from studies on *Ern2*^-/-^ mice that IRE1α cannot provide redundant functions replacing IRE1β in the development and maintenance of mucosal tissues. This is also readily apparent by analysis of their different biochemical and cellular functions. First, the endonuclease activity of IRE1α is more potent, and unlike IRE1β it requires activation by ER stress (16). Such a chronic need for stress-induced IRE1α activation to fulfill goblet cell development in the intestine and other tissues forming mucosal barriers would trigger inflammation (5, 20). And, in addition, upon induction of cell stress, we have found that IRE1β assembles with IRE1α to inhibit its activation. Thus, another part of the neofunctionalization of *Ern2* genes, we propose, was likely driven by the need to quench IRE1α hyperactivation in the context of the many, diverse, and continuous exposures to cell stress inherent to mucosal surfaces (1, 16).

How gut microbes regulate *Ern2* gene expression remains unknown. Our results suggest that specific components of the microbiota are needed to turn on goblet cell development and IRE1β function. Butyrate appears to be one component that can activate this process, although not necessarily the only component. How loss of IRE1β function enables selection of a microbial community that fails to induce IRE1β expression and function also remains unknown. A likely explanation is that *Ern2*^-/-^ mice produce altered mucins and defective mucus barriers which select for microbial species that thrive independently of IRE1β and its downstream effector functions. Our studies show, for example, that crypts from *Ern2*^-/-^ mice have reduced expression of sulfo- and sialyltransferase genes involved in mucin glycosylation. The composition of their mucus barriers likely mimics the mucin glycans produced by germ-free mice (e.g. shorter, less complex glycans) (35). Microbes that favor such a mucin environment would not benefit from induction of *Ern2* expression, which would alter goblet cell maturation and mucus assembly – thus explaining the dysbiosis and its phenotype with respect to *Ern2* expression.

Although IRE1β has been associated with increased susceptibility to intestinal inflammation and colitis in mouse models (1, 5), a definitive role for the protein in human gastrointestinal disease has not been ascribed. We note here that the colon epithelial phenotype observed in *Ern2*^-/-^ mice contains features of ulcerative colitis (UC), including defects in goblet cell maturation and mucin secretion, impaired assembly of the colon mucus layer, and dysbiosis of the gut microbiota (36–40). While the epithelium still develops as an intact monolayer in *Ern2*^-/-^ mice, the altered structure and function of the mucosal surface is associated with earlier onsets of gut infection and inflammation. Thus, it is possible the *Ern2*^-/-^ mice, and in particular the microbiota-epithelial-mucus barrier feedback loop enabled by IRE1β, may model epithelial defects associated with the onset of inflammation in UC. Although the human *ERN2* gene has not been identified as a risk allele in genome wide association studies of UC, reduced expression of *ERN2* mRNA has been found (41). Thus, stimulating IRE1β enzymatic activity directly with small molecule activators (42–45) or promoting *ERN2* expression via consortia of microbes (or their products) may restore mucus barrier function and host-microbiota crosstalk in some disease contexts.

## METHODS

### In vivo experiments in mice

All experimental procedures involving mice were approved by Boston Children’s Hospital Institutional Animal Care and Use Committee.

#### Husbandry and breeding

C57BL/6J wild type (WT) mice (#000664, The Jackson Laboratory, Bar Harbor Maine), *Ern2*^-/-^ mice (a kind gift from M. Hussain with permission from D. Ron), *Xbp1*^fl/fl^ mice (a kind gift from L. Glimcher) and Tg(*Vil1*-cre)1000Gum mice (#021504, The Jackson Laboratory, Bar Harbor, ME) were maintained in a specific pathogen-free facility under standard conditions. WT and *Ern2*^-/-^ mice were also rederived germ free and maintained in sterile isolators with autoclaved bedding, food, and water. Littermate comparisons were performed between siblings derived from *Ern2*^-/-^ and WT matings. Co-housing comparisons were performed by weaning age matched WT weanlings with *Ern2*^-/-^ weanlings from independent homozygous matings into the same cage at 3 weeks of age. Mice with Intestine-specific deletion of *Xbp1* were generated by first mating *Xbp1*^fl/fl^ mice with *Vil-Cre*^+^ mice followed by crossing of *Xbp1*^fl/+^;*Vil1-Cre*^+^ mice with *Xbp1*^fl/fl^ mice to generate *Xbp1*^fl/fl^;*Vil1-Cre*^+^ and *Xbp1*^fl/fl^;*Vil1-Cre*^-^ mice for experiments.

#### Dibenzazepine (DBZ) studies

Co-housed 8-week-old WT and *Ern2*^-/-^ mice were interperitoneally injected daily for four days with 10 μmol/kg Dibenzazepine (DBZ, previously Syncom, now Chiralix, Nijmegen, Netherlands) suspended in 0.5% (w/v) Methocel E4M (Dupont, Bellevue, WA) and 0.1% Tween 80 in water (17). Control mice were injected daily with vehicle alone (0.5% (w/v) Methocel E4M/ 0.1% Tween 80). On day 5 mice were sacrificed, a segment of distal colon was collected for histologic analysis, and the remaining distal colon was used for epithelial cell collection.

#### Citrobacter rodentium studies

Co-housed 6-8 week old WT and *Ern2*^-/-^ mice were fasted for 4 hours and then gavaged with approximately 2-4×10^8^ colony forming units (CFUs) *C. rodentium* (strain DBS100) from an overnight culture. All procedure with mice were performed in a biosafety cabinet and mice were housed in a Biosafety Level 2 room for the duration of the experiment. For quantification of *C. rodentium* in stool, fresh stool pellets were homogenized with a sterile pestle in 0.5 mL phosphate buffered saline (PBS), plated on MacConkey agar plates, and incubated overnight at 37 °C. For quantification of *C. rodentium* from colon tissue, fresh or snap frozen colon tissue segments were homogenized with a sterile pestle in PBS, plated on MacConkey agar plates, and incubated overnight at 37 °C. *C. rodentium* CFUs were determined by counting colonies from triplicate plating of a serial dilution and normalizing the number of colonies by stool or tissue weight. At defined time points during the infection or recovery after clearance, mice were sacrificed, colon tissue was collected to determine *C. rodentium* load, and a segment of distal colon was collected for histologic analysis. H&E-stained sections were evaluated by a blinded pathologist for mucosal hyperplasia, goblet cell depletion, crypt apoptosis, epithelial erosion, crypt architectural distortion, lymphocytic infiltrate intensity, neutrophilic infiltrate intensity, and submucosal inflammation and edema. Each category was assigned a score of 0-3 and summed to give a total histology score for each individual animal.

#### Colonization of germ-free mice with a gut microbiota

Germ free (GF) mice bred and raised in isolators were colonized with gut microbes derived from CONV-WT or CONV-*Ern2*^-/-^ donor mice housed in our SPF facility. Donor microbiotas were prepared by harvesting the cecum from a donor mouse, cutting the cecum open longitudinally, scraping cecal contents with forceps into a tissue grinder tube, and homogenizing in 2.5 mL sterile PBS. The cecal homogenate was centrifuged at 500xg for 3 min, and the supernatant was immediately used for colonization into recipient mice. Recipient mice were colonized by one-time gavage with 200 μL of cecal homogenate and housed for the duration of the experiment in sterile cages outside of the germ-free isolator. At 14 days post-colonization, mice were sacrificed, and a segment of distal colon was collected for histologic analysis, and the remaining distal colon was used to isolate colon crypt epithelial cells.

#### Colonization of germ-free mice followed by TUDCA administration

WT and *Ern2*^-/-^ germ free mice were colonized with a microbiota from WT donor mice as described above. Beginning the day after colonization, mice were gavaged every other day for 12 days with 500 mg/kg sodium tauroursodeoxycholate in PBS (TUDCA, Sigma Aldrich, St. Louis, MO) (22). At 14 days post-colonization, mice were sacrificed, and a segment of distal colon was collected for histologic analysis.

#### Antibiotic treatment followed by co-housing to recolonize gut microbes

WT female mice were treated with an antibiotic cocktail (ampicillin (0.5 g/L), vancomycin (0.25 g/L), metronidazole (0.5 g/L), and neomycin (0.25 g/L) (Sigma Aldrich, St. Louis, MO) dissolved in drinking water) for 7 days (46). After 7 days, antibiotic-treated mice were recolonized by co-housing with conventionally raised WT or *Ern2*^-/-^ donor mice for 14 days. At 14 days post-colonization, mice were sacrificed, and a segment of distal colon was collected for histologic analysis. Colon tissue was also collected from age-matched mice that did not receive antibiotic and mice that received antibiotics for 7 days but without subsequent co-housing.

#### Sodium butyrate administration in germ free mice

WT and *Ern2*^-/-^ germ free mice received either regular drinking water or drinking water supplemented with 100 mM sodium butyrate (#A11079 Alfa Aesar, Tewksbury, MA) (47) for 14 days. Mice were monitored for normal drinking behavior. After 14 days, mice were sacrificed, and a segment of distal colon was collected for histologic analysis.

#### Isolation of colon epithelial cells and intact colon crypts

Colon tissue was harvested, lumenal contents were gently removed, and the tissue was flushed with ice-cold PBS. The tissue was cut open longitudinally, cut into small pieces, washed 3 times with ice cold PBS, and incubated in PBS with 10 mM EDTA for 45 min at 4 °C on a rotary shaker. Epithelial cells were dissociated by vigorous shaking for 5 min, and the cell suspension was decanted into base media (Advanced DMEM/F12 Media, 20% FBS, 10 mM HEPES, 100 μg/mL Penicillin/Streptomycin, 2 mM L-glutamax). The remaining tissue pieces were subjected to a second round of epithelial cell isolation in the same manner. To isolate intact colon crypts, the epithelial cell suspension was passed through a 100 μm strainer followed by a 40 μm strainer. Intact crypts retained on the 40 μm strainer were washed, eluted with base media, and pelleted by centrifugation at 300 xg for 3 min. Crypts were either used to generate organoid lines as described below, or washed with PBS and stored at −80 °C for later use.

### In vitro experiments with colonoids and cell lines

#### Preparation of epithelial colonoid lines

Three independent epithelial colonoid lines were prepared from WT and *Ern2*^-/-^ mice. Intact colon crypts were harvested from mice as described above. Crypts were resuspended in Matrigel (Corning) on ice and 30 μL drops were plated in 24-well plates (typically 3-4 drops for the initial isolation) and cultured in Complete media (base media + 50% WRN-conditioned media prepared from L cells expressing Wnt/R-spondin/Noggin as described (48)) with 10 μM Y27632 (ROCK inhibitor, Sigma-Aldrich Y0503) at 37 °C 5% CO2. Media was changed every other day. Cultures were passaged by dissolving Matrigel with Cell Recovery Solution, washing colonoids with base media, mechanically disrupting colonoids by pipetting, and plating in 1.5- to 2-fold more Matrigel than used in the previous plating. Cultures were expanded as needed for experiments and cryopreservation of colonoid lines.

#### DAPT-induced differentiation assay

For differentiation experiments, WT and *Ern2*^-/-^ colonoids were plated in Matrigel and cultured in Complete media supplemented with 10 mM Y27632 for 24 hr. After 24 hr, colonoids were treated with 10 μM DAPT (Sigma-Aldrich 565770) ± 50 μM 4μ8C (Sigma-Aldrich 412512) for an additional 24 hr; control colonoids had Complete media alone or Complete media with 50 μM 4μ8C (no DAPT). Bright field images of colonoids were collected at three different focal planes over the Matrigel drop using a Cytation 5 imager (BioTek). Colonoids that were in focus were scored as either spheroid or non-spheroid, and the percentage of colonoids with a spheroid morphology was calculated for each condition and fractional change in the percent spheroid morphology was expressed relative to the control colonoids (no DAPT, no 4μ8C). The experiment was performed on 2-3 independent colonoid lines in at least in 2 independent experiments for each line. Colonoids were collected for RNA extraction and expression analysis of goblet cell marker genes by qPCR.

#### Butyrate-induced Ern2 expression in epithelial cell lines

T84 cells were maintained in 1:1 DMEM/F12 media supplemented with 6% newborn calf serum. Cells were plated on 0.33 cm^2^ Transwell inserts with 3 μm pore size polyester membranes and allowed to polarize for 7 days. Monolayer formation was assessed by measuring transepithelial electrical resistance with an epithelial volt/Ohm meter (EVOM; World Precision Instruments). Monolayers were treated with 10 mM sodium butyrate, 10 mM sodium acetate, or media alone for 24 hrs. LS174T cells were cultured in DMEM supplemented with 10% FBS and 1x non-essential ammino acid solution (Gibco). Cells were treated with 1 mM sodium butyrate or control media for 24 hr. After cell treatments, cells were washed with PBS and RNA was extracted for expression analysis.

#### IRE1β Xbp1 splicing activity in HEK293^doxIRE1β^ cell line

HEK293^doxIRE1β^ cell line was described by Grey, Cloots, Simpson, et al. (16). Expression of Flag-tagged human IRE1β in this cell line was induced by treatment with 0, 10, or 100 ng/mL doxycycline for 24 hr. RNA was extracted for expression analysis by qPCR (spliced Xbp1 transcript) and RNAseq as described below.

### Histologic analysis of mouse colon tissues

#### Histologic analysis of colon crypt length and goblet cells

A segment (1 cm) of distal colon was isolated, lumenal content was gently removed, and the tissue was fixed in 10% formalin for 24-48 hr at room temperature. Fixed tissue was embedded in paraffin and sections prepared for histologic staining with H&E or Alcian blue. Sections were also stained with anti-Ki67 (Cell Signaling Technologies, CST12202S, 1:500 dilution) and anti-MUC2 (Santa Cruz Biotechnology, sc-15334, 1:100 dilution) antibodies and Cy3-labeled anti-rabbit secondary antibodies. Images of stained sections were collected on an Olympus BX41 microscope with a Pixelink PL-D682CU color CMOS camera for brightfield and epifluorescence imaging. Crypt length was measured in H&E- or AB-stained sections in well-oriented crypts that extend the full-length of the mucosa. Goblet cells were enumerated by counting AB^+^, PAS^+^, or anti-MUC2^+^ mucin granules in the upper half of well-defined crypts (measured from the lumenal opening of the crypt toward the base) and normalizing by the total number of IECs (stained nuclei) lining the crypt. In most cases, crypt length and goblet cell numbers were averaged over at least 5 well-defined crypts. Goblet cell theca area was measured in ImageJ by measuring the AB-stained area for goblet cells in the upper half of well-oriented crypts.

#### Histologic analysis of colon mucus layer

To preserve the colon mucus layer for analysis, a 2-3 cm segment of distal colon containing a fecal pellet was excised and fixed in Carnoy’s solution (60% methanol, 30% chloroform, 10% acetic acid) for 24-48 hr at room temperature (49). Fixed tissue was washed twice with 100% methanol followed by two washes with 100% ethanol. The fixed tissue segment was cut into pieces and embedded in paraffin for cross-section cuts through the fecal pellet. No water was used in any of the fixation or processing steps to ensure that the mucus layer was preserved. Slides were stained with AB/PAS. The inner mucus layer was analyzed by measuring thickness of AB/PAS-stained region adjacent to the epithelium at regularly spaced intervals in images covering the entire section around a fecal pellet (usually in 2-3 slices of a single fecal pellet) and averaged over the entire distribution of measurements for an individual animal. Carnoy’s-fixed tissue was also stained with the in situ hybridization probe EUB338 to detect lumenal bacteria as described (49). Slides were deparaffinized, stained with 20 mg/mL of 594-EUB338 probe (AlexaFluor594-5’-GCTGCCTCCCGTAGGAGT -3’, Integrated DNA Technologies) or 594-CONTROL probe (AlexaFluor594-5’-CGACGGAGGGCATCCTCA-3’, Integrated DNA Technologies) in hybridization buffer (20 mM Tris pH 7.4, 0.9 M NaCl, 0.1% (w/v) SDS) at 50 °C overnight in a humidified chamber. After staining, slides were incubated in FISH washing buffer (20 mM Tris pH 7.4, 0.9 M NaCl) for 20 min at 50 °C then washed three times with phosphate buffered saline. Sections were co-stained with DAPI (D3571, Thermo Fisher) and slides were mounted using Prolong Antifade (P36961, Thermo Fisher).

### Molecular analysis of mRNA gene expression

#### Expression analysis by RNAseq

RNA was extracted from intact colon crypts and cell lines using the RNeasy Mini extraction kit (Qiagen). For RNA sequencing, library preparation, next generation sequencing, and data processing was performed by the Dana Farber Cancer Institute Molecular Biology Core Facility. Total RNA quality and concentration were measured using Aligent Bioanalyzer and RNA libraries were prepared using KAPA stranded mRNA HyperPrep kits. RNA was sequenced by synthesis with 75 cycles of single end reads on an Illumina NextSeq500. Data were processed using the Visualization Pipeline for RNAseq (VIPER) (50). Sequence reads were aligned using STAR (51) and raw gene counts were used to calculate differential expression (log_2_ [Group1/Group2]) between groups with DESeq2 (52). Genes with genome-wide adjusted *p_adj_* < 0.01 were considered significantly different. Gene set enrichment analysis was performed by calculating hypergeometric distributions for differentially expressed genes (*p_adj_* < 0.01) using epithelial cell signatures derived from Haber, et. al. (2). Functional analysis of differentially expressed genes was performed using gProfiler (53).

#### Expression analysis by qPCR

RNA was extracted from intact colon crypts, colon epithelial cells, colonoids, and cell lines using the RNeasy Mini extraction kit (Qiagen). cDNA was prepared from total RNA using the iScript cDNA Synthesis Kit (BioRad). Target gene cDNA was amplified using primers (Table S1) and Sso Advanced Universal SYBR Green Supermix (BioRad) on a CFX384 Touch Real-Time PCR Detection system (BioRad). Reactions were assayed in triplicate for each sample, and the average Cq value was used to calculate the mean expression ratio of the test sample compared to the control sample using the 2-ΔΔCt method. Cq values for targets were analyzed relative to Cq values for *Hprt* and *Ppia* reference genes.

### Molecular and biochemical analysis of mouse gut microbiome

#### Metagenomic sequencing

Stool pellets from co-housed WT and *Ern2*^-/-^ mice were collected directly into sterile tubes, frozen on dry ice, and stored at −80 °C. DNA was extracted using ZymoBIOMICS DNA miniprep kit (D4300, Zymo Research) and metagenomic DNA sequencing libraries were constructed using the Nextera XT DNA Library Prep Kit (FC-131-1096, Illumina) and sequenced on a NextSeq500 Sequencing System as 2 x 150 nucleotide paired-end reads. Shotgun metagenomic reads were first trimmed and quality filtered to remove sequencing adapters and host contamination using Trimmomatic (54) and Bowtie2 (55), respectively, as part of the KneadData pipeline (https://bitbucket.org/biobakery/kneaddata). Metagenomic data was profiled for microbial taxonomic abundances and microbial metabolic pathways using Metaphlan3 (56) and HUMAnN3 (57), respectively.

#### Short chain fatty acid analysis

Stool pellets from WT and *Ern2*^-/-^ mice were collected directly into sterile pre-weighed tubes, immediately frozen, and stored at −80 °C. Pellets were homogenized in HPLC grade water by vortexing. The pH of the cleared homogenate was adjusted to 2-3, 2-methyl pentanoic acid was added as an internal standard (0.1%), and SCFAs were extracted by adding 1 volume of ethyl ether anhydrous. Samples were vortexed for 2 min and centrifuged at 5000 g for 2 min. The upper ether layer was collected and SCFA content was analyzed on an Agilent 7890B gas chromatography system with flame ionization detector using a high-resolution capillary column for detection of volatile acids (DB-FFAP, 30 m x 0.25 mm with 0.25 μm film thickness; Agilent Technologies, Santa Clara, CA). A standard solution containing 10 mM of acetic, propionic, isobutyric, butyric, isovaleric, valeric, isocaproic, caproic, and heptanoic acids (Supelco CRM46975, Bellefonte, PA) was processed and analyzed in the same manner as the stool samples. The retention times and peak heights of the acids in the standard mix were used as references for the unknown samples. Each acid was identified by its specific retention time and the concentration was determined and expressed as mM per gram of fecal material. Chromatograms and data integration were carried out using OpenLab ChemStation Software (Agilent Technologies).

## ACKNOWLEDGEMENTS

We thank members of the Lencer lab for valuable discussions throughout the course of this project; David Ron for providing *Ern2*^-/-^ mice and Laurie Glimcher for providing *Xbp1*^fl/fl^ mice used in these studies; the Harvard Digestive Disease Center for services provided through the Microscopy and Histopathology Core, the Epithelial Cell and Mucosal Immunology Core, and the Gnotobiotic, Microbiology, and Metagenomics Core; and the Molecular Biology Core Facility at the Dana Farber Cancer Institute for RNA sequencing. This work was supported by National Institutes of Health grants K01DK119414 (M.J.G.), R01DK048106 (W.I.L. and M.J.G.), R01DK125407 (B.A.M.), R01DK061931 (J.R.Turner), and P30DK034854 (W.I.L.).

## AUTHOR CONTRIBUTIONS

M.J.G. and W.I.L. conceived the project. M.J.G., H.D., D.V.W., S.E.F., J.R.Thiagarajah., B.A.M., J.R.Turner., and W.I.L. contributed to the experimental design. M.J.G., H.D., D.V.W., and I.A.M.K. conducted experiments and processed data. M.J.G., H.D., J.R.Turner., and W.I.L. analyzed data and interpreted results. M.J.G., H.D., and W.I.L. wrote the manuscript. All authors reviewed the manuscript prior to submission.

## CONFLICT OF INTEREST

J. R. Turner is a founder and shareholder of Thelium Therapeutics and has served as a consultant for Entrinsic, Immunic, Johnson & Johnson, Kallyope, and 89Bio.

---

**Supplemental Figure S1.**
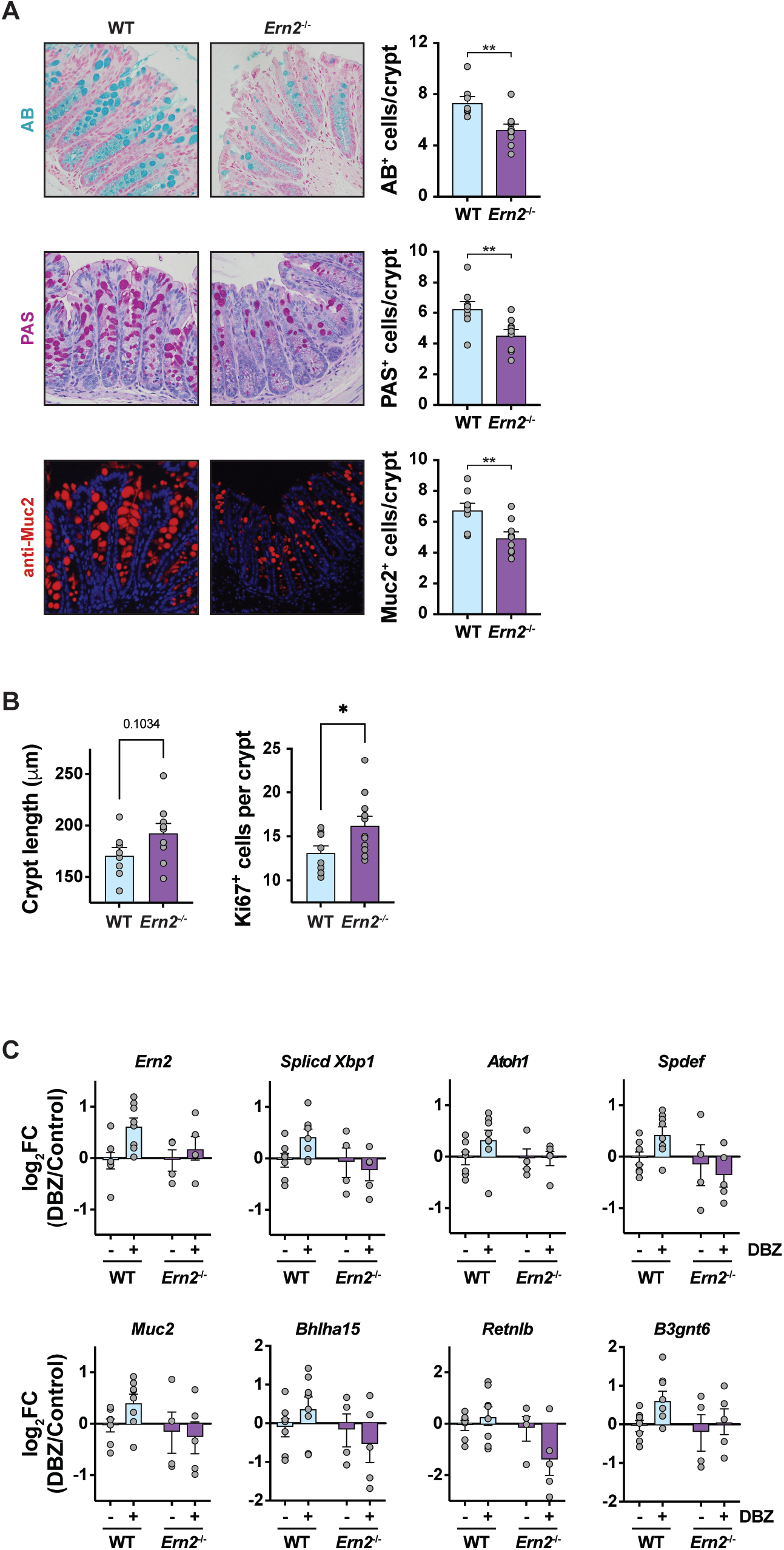
**(supporting Figure 1).** (A) Representative images of Alcian blue (AB), Periodic acid Schiff (PAS) and anti-MUC2 antibody-stained sections of distal colon from CONV-WT and CONV-*Ern2*^-/-^ mice (littermates). Bar graphs show the number of AB+, PAS+, and anti-MUC2^+^ cells in the upper half of well-defined crypts. Symbols represent the average value for an individual mouse. Bars represent mean ± SEM. Mean values were compared by unpaired t-test. (B) Bar graphs show crypt length and the number of anti-Ki67^+^ cells measured in well-oriented crypts in the distal colon. Symbols represent the average value for an individual mouse. Bars represent mean ± SEM. Mean values were compared by unpaired t-test. (C) Bar graphs show relative mRNA expression levels of goblet cell marker genes measured by qPCR in colon epithelial cells isolated from CONV-WT and CONV-*Ern2*^-/-^ mice treated with and without DBZ. Symbols represent relative expression for individual mice and bars represent mean ± SEM. These data are summarized as a heat map in Fig. 1D.

**Supplemental Figure S3.**
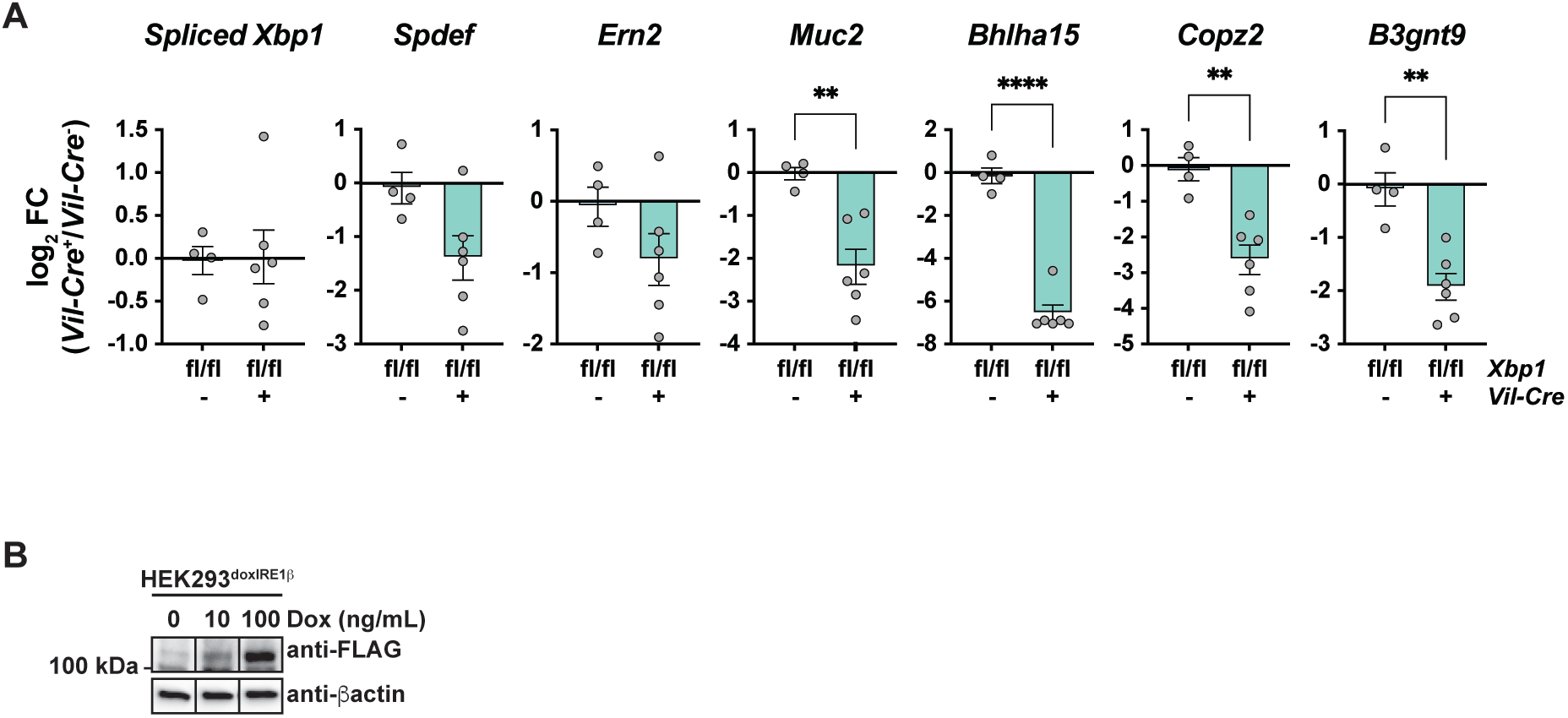
**(supporting Fig. 3)** (A) Bar graphs show relative mRNA expression levels of goblet cell marker genes measured by qPCR in colon crypt epithelial cells isolated from CONV-*Xbp1*^fl/fl^;*Vil-Cre*^-^ and CONV-*Xbp1*^fl/fl^;*Vil-Cre*^+^ mice. Symbols represent relative expression for individual mice and bars represent mean ± SEM. These data are summarized as a heat map in Fig. 3B. (B) Representative western blot showing doxycycline-induced expression of FLAG-tagged IRE1β in HEK293^doxIRE1β^ cell line. Cells were treated with indicated contraction of Dox for 20 hrs to induce expression. The indicated slices all come from the same gel and membrane that was cropped to exclude intervening lanes that contain other treatment conditions not considered here. Data are reproduced from (16) in support of IRE1β-induced *Xbp1* mRNA splicing, XBP1-dependent gene expression, and ER/secretory compartment gene expression shown in Fig. 3D.

